# Metatranscriptomics as a tool to identify fungal species and subspecies in mixed communities

**DOI:** 10.1101/584649

**Authors:** Vanesa R. Marcelino, Laszlo Irinyi, John-Sebastian Eden, Wieland Meyer, Edward C. Holmes, Tania C. Sorrell

## Abstract

High-throughput sequencing (HTS) enables the generation of large amounts of genome sequence data at a reasonable cost. Organisms in mixed microbial communities can now be sequenced and identified in a culture-independent way, usually using amplicon sequencing of a DNA barcode. Bulk RNA-seq (metatranscriptomics) has several advantages over DNA-based amplicon sequencing: it is less susceptible to amplification biases, it captures only living organisms, and it enables a larger set of genes to be used for taxonomic identification. Using a defined mock community comprised of 17 fungal isolates, we evaluated whether metatranscriptomics can accurately identify fungal species and subspecies in mixed communities. Overall, 72.9% of the RNA transcripts were classified, from which the vast majority (99.5%) were correctly identified at the species-level. Of the 15 species sequenced, 13 were retrieved and identified correctly. We also detected strain-level variation within the *Cryptococcus* species complexes: 99.3% of transcripts assigned to *Cryptococcus* were classified as one of the four strains used in the mock community. Laboratory contaminants and/or misclassifications were diverse but represented only 0.44% of the transcripts. Hence, these results show that it is possible to obtain accurate species- and strain-level fungal identification from metatranscriptome data as long as taxa identified at low abundance are discarded to avoid false-positives derived from contamination or misclassifications. This study therefore establishes a base-line for the application of metatranscriptomics in clinical mycology and ecological studies.

## Introduction

Microscopic fungal species, such as yeasts and some filamentous fungi, do not live in isolation, they are most commonly found within mixed microbial communities inhabiting soil, water systems, plants and animal hosts. Assessing the diversity of fungi in mixed communities is important because different fungal taxa may exhibit distinctive phenotypes, and consequently may have different pathogenicity or functional roles. For example, in the rhizosphere, changes in fungal community composition have been associated with shifts in nutrient cycling (Hannula *et al.* 2017). Humans also harbor, or are exposed to, a diverse fungal community that provides a source of opportunistic pathogens (Bandara *et al.* 2019; Huffnagle & Noverr 2013; Seed 2014). Although it is typically assumed that invasive fungal infections are caused by a single strain, multiple *Candida* strains have been observed during the course of a single episode of infection (Soll *et al.* 1988). Furthermore, nearly 20% of patients with cryptococcosis are infected with multiple strains, with different phenotypes and virulence traits (Desnos-Ollivier *et al.* 2015; Desnos-Ollivier *et al.* 2010). Strain-level fungal diversity may influence therapeutic responsiveness and needs further investigation.

Despite its importance, fungal taxonomic diversity is poorly characterized. From over two million fungal species estimated to exist, less than 8% have been described (Hawksworth & Lucking 2017). Even well-known fungal species are often overlooked during routine diagnostic procedures, surveillance and biodiversity surveys (Brown *et al.* 2012; Enaud *et al.* 2018; Yahr *et al.* 2016). This is in part due to challenges in the detection and classification of these organisms, especially microscopic and cryptic species, for example, the etiologic agents of cryptococcosis. Currently, two species complexes are recognized: *Cryptococcus neoformans* and *Cryptococcus gattii* (Kwon-Chung *et al.* 2002). Seven major haploid lineages are found within these two species complexes (*C. neoformans* species complex: VNI, VNII, VNIV, and *C. gattii* species complex: VGI, VGII, VGIII and VGIV) and their recognition as distinct biological species has been debated (Hagen *et al.* 2015; Kwon-Chung *et al.* 2017; Ngamskulrungroj *et al.* 2009). Being able to distinguish closely-related lineages is important because their phenotype, virulence and ecophysiology can vary substantially. For example, the JEC21 and B-3501 strains of *C. neoformans* var. *neoformans* (VNIV) are 99.5% identical at the genomic sequence level but differ substantially in thermotolerance and virulence (Loftus *et al.* 2005). Likewise, different virulence and antifungal tolerance traits were observed within lineages of *C. gattii* VGIII (Firacative *et al.* 2016).

The introduction of high-throughput sequencing (HTS) marked a new era in mycological research, where the vast diversity of fungi can be studied without the need for culture (Nilsson *et al.* 2019). To date, amplicon sequencing of marker genes (metabarcoding) has been the most popular HTS method used to identify fungal species in mixed communities. Despite its indisputable utility, metabarcoding surveys are affected by PCR amplification biases, and even abundant species can go undetected due to primer mismatch (Marcelino & Verbruggen 2016; Nilsson *et al.* 2019; Tedersoo *et al.* 2015). In addition, DNA fragments from dead organisms inflate biodiversity estimates in metabarcoding surveys (Carini *et al.* 2016). Stool samples, for instance, naturally contain food-derived DNA, which cannot be distinguished from the genetic material of the resident gut microbiota when using DNA-based methods. These challenges can be circumvented by directly sequencing actively transcribed genes, via RNA-Seq, hence avoiding the amplification step, and obtaining an unbiased characterization of the living microbial community. Metatranscriptomics has been used to identify RNA viruses in a range of animal samples (Shi *et al.* 2016; Shi *et al.* 2017; Wille *et al.* 2018; Zhang *et al.* 2018) and to characterize the functional profile of microbial communities (Bashiardes *et al.* 2016; Kuske *et al.* 2015). Studies applying metatranscriptomics to mycorrhizal communities have provided valuable insights into the functional roles of fungi in these symbiotic systems (Gonzalez *et al.* 2018; Liao *et al.* 2014). However, links between functional and species-level taxonomy have been sought infrequently, likely because fungal identification from metatranscriptome data is considered unreliable below phylum level (Nilsson *et al.* 2019). Critically, it is currently unknown whether metatranscriptomics can accurately identify fungi at the species and subspecies level within a mixed community. This information is fundamental to the investigation of the potential and utility of metatranscriptomics in diagnostics and ecological studies.

Herein, we evaluated the utility of metatranscriptomics as a tool for the simultaneous identification of fungal species, using a defined mock community containing 15 fungal species. In addition, we investigated whether strains belonging to sister species, such as the *C. neoformans* and *C. gattii* species complexes could be identified correctly using metatranscriptomics. Rather than focusing on marker genes, we sought to classify fungal species using the information from all expressed genes, using the totality of NCBI’s nucleotide collection as a reference database. This study paves the way to apply state-of-the art techniques in fungal biodiversity surveys and clinical diagnostics.

## Methods

A defined fungal community was constructed from 17 isolates, including 15 fungal species and three strains of the *C. neoformans* species complex in addition to one strain of *C. gattii* (Table 1). Fungal strains were obtained from the Westmead Mycology Culture Collection and cultured on Sabouraud agar at 27°C for 72 hours. A sweep of colonies was made with a disposable inoculating loop and dispersed in PBS. Fungal cells were quantified in a Neubauer chamber and their concentration adjusted such that the fungal mixture contained equal concentrations of each species (10^8^ cells/mL). RNA was isolated with the RNeasy Plus kit (Qiagen), following the manufacture’s protocol, with an initial freeze-thaw step in liquid nitrogen to disrupt fungal cells. The quantity and quality of the RNA extract was determined with the Nanodrop Spectrophotometer (Thermo Scientific) and the Agilent 2200 TapeStation. As some residual DNA was detected, the RNA extract was further treated with DNase I (Qiagen). Ribosomal depletion (Ribo-Zero Gold technology), library preparation and sequencing (Illumina HiSeq HT, 125bp Paired End) were performed by the Australian Genomics Research Facility. The raw sequence data were deposited in the NCBI Short Read Archive (accession PRJNA521097).

**Table 1.**
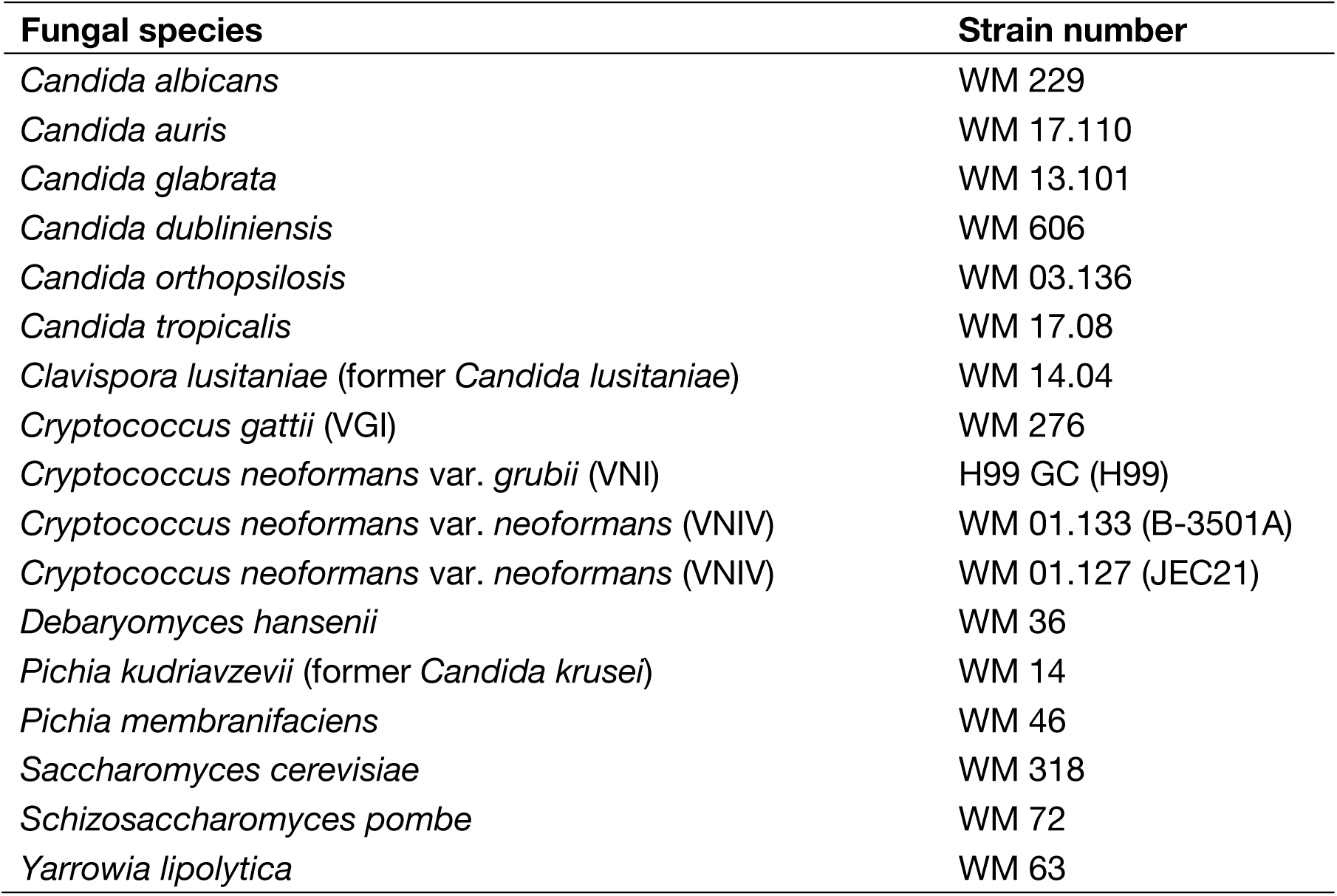
Species and strains used to construct a mock fungal community for metatranscriptome sequencing.

| Fungal species | Strain number |
| --- | --- |
| *Candida albicans* | WM 229 |
| *Candida auris* | WM 17.110 |
| *Candida glabrata* | WM 13.101 |
| *Candida dubliniensis* | WM 606 |
| *Candida orthopsilosis* | WM 03.136 |
| *Candida tropicalis* | WM 17.08 |
| *Clavispora lusitaniae* (former *Candida lusitaniae*) | WM 14.04 |
| *Cryptococcus gattii* (VGI) | WM 276 |
| *Cryptococcus neoformans* var. *grubii* (VNI) | H99 GC (H99) |
| *Cryptococcus neoformans* var. *neoformans* (VNIV) | WM 01.133 (B-3501A) |
| *Cryptococcus neoformans* var. *neoformans* (VNIV) | WM 01.127 (JEC21) |
| *Debaryomyces hansenii* | WM 36 |
| *Pichia kudriavzevii* (former *Candida krusei*) | WM 14 |
| *Pichia membranifaciens* | WM 46 |
| *Saccharomyces cerevisiae* | WM 318 |
| *Schizosaccharomyces pombe* | WM 72 |
| *Yarrowia lipolytica* | WM 63 |

Sequence reads containing more than five ambiguous positions or with average quality scores ≤ 25 were filtered from the dataset using prinseq-lite v.0.20.4 (Schmieder & Edwards 2011) with the options -ns_max_n 5 -min_qual_mean 25 -out_format 3. Assembly of sequence reads into contigs was performed with Trinity v.2.5.1 (Grabherr *et al.* 2011). Contigs were mapped to the NCBI nucleotide collection using KMA (Clausen *et al.* 2018), a novel approach that has proven to be more accurate than other mapping software. Prior to mapping, NCBI’s taxonomic identifier codes (taxids) were appended to each sequence record in the nucleotide collection, and the reference database was indexed using KMA’s options -NI -Sparse TG. Contigs were then mapped to the indexed database with the options -mem_mode -and -apm f. Matches to the reference database with low support (*i.e.* coverage < 20 and depth < 0.05) were excluded from the analyses. The species-level taxonomic classifications were based on NCBI’s taxonomy identifiers (taxids) to minimize the issue of changing species nomenclature (Federhen 2012). Subspecies-level classifications within the *Cryptococcus neoformans* and *C. gattii* species complexes were examined manually.

Abundance was estimated at the level of sequence reads and transcripts. For read-level abundances, sequence reads were mapped to transcripts using Bowtie2 (Langmead & Salzberg 2012) and quantified in Transcripts Per Million (TPM) with RSEM (Li & Dewey 2011), using the Trinity pipeline. For transcript-level abundances, the depth values estimated within KMA were used, which is the total number of nucleotides (in each contig) covering the reference sequence divided by the length of the reference sequence. The number and length of assembled contigs for each taxon is likely a better proxy for species abundance than read-level abundances (which are subject to gene expression), and therefore were used for graphic representation and analyses. For simplicity, we refer as ‘abundance’ the transcript-level abundance, unless otherwise stated.

It would be logical to expect that species with larger and gene-rich genomes would express a greater number of transcripts. To test for this potential correlation, genome sizes and the estimated number of proteins were obtained from the Fungal Genome Size Database (Kullman *et al.* 2005), Loftus et al. (2005), Muñoz et al. (2018) and NCBI’s Genome database (Supplementary table S1). The correlation coefficients between genome size, number of proteins and abundance of transcripts were estimated using Person’s correlation and visualized using the R package *ggpubr* (Kassambara 2017).

## Results

RNA sequencing yielded a total of 26,558,491 paired end reads, of which 98.3% passed quality control. Overall, 277,404 contigs (transcripts) were obtained, from which 202,219 (72.9%) were classified. The majority of the sequence reads (80.2%) mapped to a classified contig. Of the 15 fungal species sequenced, 13 were retrieved and correctly classified at the species level (Figure 1, Table 2, Supplementary table S2). The two false-negatives were *Debaryomyces hansenii* and *Schizosaccharomyces pombe*; these may have been misclassified as another fungus or were lost due to cell pooling inaccuracy and/or RNA extraction biases. A small proportion of bacterial transcripts (0.03%) and other eukaryotic microbes (0.4%, including 31 fungi that were not present in the mock community) was also observed (Table 2, Supplementary table S2), which likely represent laboratory contaminants and misclassifications (see discussion). However, these were present at a consistently lower frequency than true members of the mock community, with the most common – *Candida glycerinogenes* – only present in 0.08% of the transcripts. Some of the transcripts were assigned to entries in GenBank that do not have a species-level classification (*e.g. Candida* sp. and *Pichia* sp.). These assignments were considered misclassifications here, although it is possible that the species in our mock community are the correct species-level identity of these GenBank sequences.

**Table 2.** Abundance of reads (TPM) and abundance of transcripts (Depth) per fungal species detected with metatranscriptomics. True members of the mock community – at species level – are shown in bold.

| Species* | TPM<br>(read-level) | Depth<br>(transcript-level) | Relative abundance<br>(transcript-level %) |
| --- | --- | --- | --- |
| ***Cryptococcus neoformans*** | **149692.28** | **11049.04** | **22.464** |
| ***Candida tropicalis*** | **142496.47** | **9424.30** | **19.161** |
| ***Pichia kudriavzevii*** | **62133.74** | **9234.05** | **18.774** |
| ***Clavispora lusitaniae*** | **57317.81** | **7107.80** | **14.451** |
| ***Candida auris*** | **13402.41** | **3354.39** | **6.820** |
| ***Candida dubliniensis*** | **52027.94** | **1706.41** | **3.469** |
| ***Pichia membranifaciens*** | **10860.75** | **1498.26** | **3.046** |
| ***Candida albicans*** | **42948.69** | **1441.56** | **2.931** |
| ***Yarrowia lipolytica*** | **52376.03** | **1384.29** | **2.814** |
| ***Candida orthopsilosis*** | **44531.25** | **1308.09** | **2.660** |
| ***Candida glabrata*** | **50404.11** | **992.55** | **2.018** |
| ***Cryptococcus gattii* VGI** | **9350.81** | **463.71** | **0.943** |
| *Candida glycerinogenes* | 2345.13 | 41.51 | 0.084 |
| *Nakaseomyces delphensis* | 12821.76 | 35.45 | 0.072 |
| *Candida parapsilosis* | 15989.78 | 31.11 | 0.063 |
| *Candida nivariensis* | 1411.36 | 12.72 | 0.026 |
| *Kluyveromyces marxianus* | 47.70 | 8.90 | 0.018 |
| *Torulaspora delbrueckii* | 15.01 | 6.00 | 0.012 |
| *Kluyveromyces lactis* | 9.24 | 3.93 | 0.008 |
| ***Saccharomyces cerevisiae*** | **22.84** | **3.77** | **0.008** |
| *Eremothecium sinecaudum* | 25.93 | 3.67 | 0.008 |
| *Pichia cecembensis* | 751.23 | 3.60 | 0.007 |
| *Lodderomyces elongisporus* | 34.30 | 3.59 | 0.007 |
| Uncultured *Candida* | 885.35 | 3.13 | 0.006 |
| *Eremothecium gossypii* | 10.13 | 3.04 | 0.006 |
| *Naumovozya dairenensis* | 17.37 | 2.80 | 0.006 |
| *Suhyomyces tanzawaensis* | 30.87 | 2.25 | 0.005 |
| Dipodascaceae sp. LM136 | 24286.11 | 2.16 | 0.004 |
| *Cyberlindnera jadinii* | 12.32 | 2.04 | 0.004 |
| *Metschnikowia bicuspidata* | 16.02 | 1.29 | 0.003 |
| *Brettanomyces naardenensis* | 96.13 | 1.19 | 0.002 |
| *Pichia norvegensis* | 783.06 | 1.11 | 0.002 |
| *Debaryomyces fabryi* | 22.87 | 0.92 | 0.002 |
| *Candida neerlandica* | 487.65 | 0.69 | 0.001 |
| *Melanotaenium endogenum* | 262.12 | 0.59 | 0.001 |
| *Pichia kluyveri* | 51814.00 | 0.55 | 0.001 |
| *Candida pseudohaemulonis* | 560.14 | 0.49 | 0.001 |
| *Candida* sp. (in: *Saccharomycetales*) | 330.15 | 0.49 | 0.001 |
| *Pichia* sp. 2 TMS-2011 | 0.00 | 0.45 | 0.001 |
| *Cryptococcus neoformans* AD hybrid | 0.00 | 0.44 | 0.001 |
| *Saccharomycetales* sp. LM594 | 2.60 | 0.30 | 0.001 |
| *Naumovozya castellii* | 4.56 | 0.29 | 0.001 |
| *Saccharomyces pastorianus* | 0.81 | 0.26 | 0.001 |
| *Cryptococcus gattii* VGIII | 0.48 | 0.21 | 0.000 |
| Other Eukaryotes | 936.75 | 18.83 | 0.038 |
| Bacteria | 423.36 | 15.79 | 0.032 |
| Unclassified | 198000.57 | 8.00 | 0.016 |
| **TOTAL** | **1000000** | **49186** | **100** |

\* Species defined according to the NCBI taxonomy database. Strain numbers may indicate vouchers rather than genetically different lineages.

**Figure 1.**
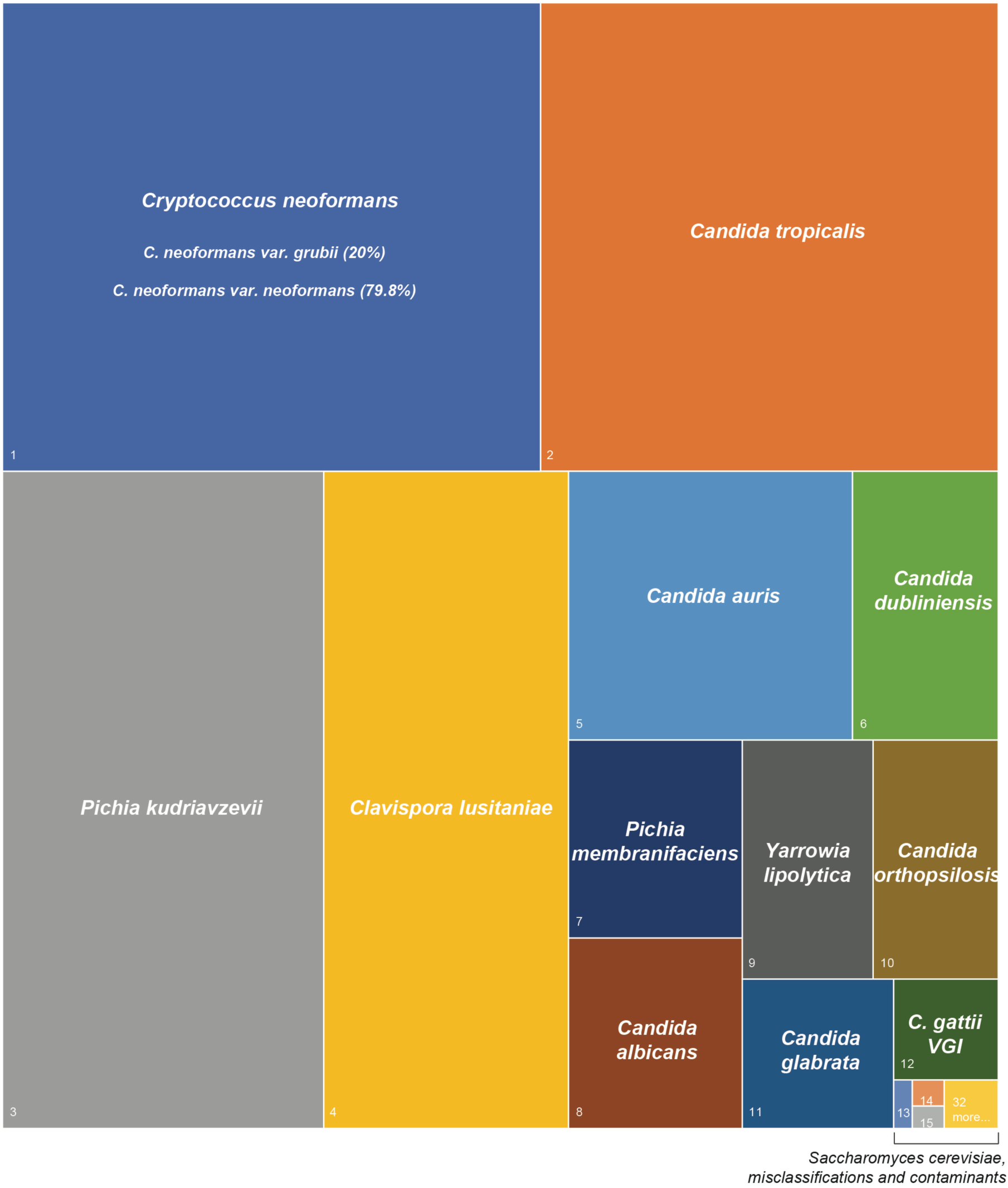
Relative abundance of transcripts assigned to microbial species recovered in the metatranscriptome of a mock community. See Table 2 for a full list of species and more details about their abundance.

Overall, the commonest species detected was *C. neoformans*, which was to be expected as it comprised three strains in the mock community and therefore was three times more abundant than other fungal species. Transcripts belonging to *Candida tropicalis* and *Pichia kudriavzevii* (former *Candida krusei*) – were also common (19.2% and 18.8%, respectively), while *C. albicans, C. orthopsilosis* and *C. glabrata* (other causes of candidaemia in humans) were detected at lower abundance (2.0 – 2.9%). There was no relationship evident between abundance of transcripts and phylogenetic relatedness. Genomes with low GC content can be overrepresented in metagenomic sequencing (Shakya *et al.* 2013). Conversely, some of the species detected here in high abundance (*Cryptococcus neoformans* and *Clavispora lusitaniae*) have a higher GC content than most other fungal species (Dujon 2010), suggesting that GC bias is unlikely to affect our results. No correlation between abundance of transcripts and genome size or estimated number of proteins was observed (*p* > 0.05, Supplementary figure 1).

Molecular type and strain-level variation within the *Cryptococcus neoformans* and *C. gattii* species complexes was also detected, with contigs matching to *C. gattii* VGI WM 276, *C. neoformans* var. *grubii* VNI H99 and *C. neoformans* var. *neoformans* VNIV strains B-3501A and JEC21 (Figure 2, Supplementary table S3). A proportion of the transcripts (1.6%) matched with equal probability scores to both strains of *C. neoformans* var. *neoformans* (B-3501A and JEC21, Supplementary tables S2 and S3). From the transcripts classified as *Cryptococcus* spp., 99.3% were classified as one of the four *Cryptococcus* strains (or both B-3501A and JEC21) used in the mock community. It is possible that misclassifications occurred within the strains analyzed. For example, transcripts originally from JEC21 might have been classified as B-3501A and *vice versa*. As it is not possible to know from which strain the transcripts originated, these possible misclassifications would be undetected.

**Figure 2.**
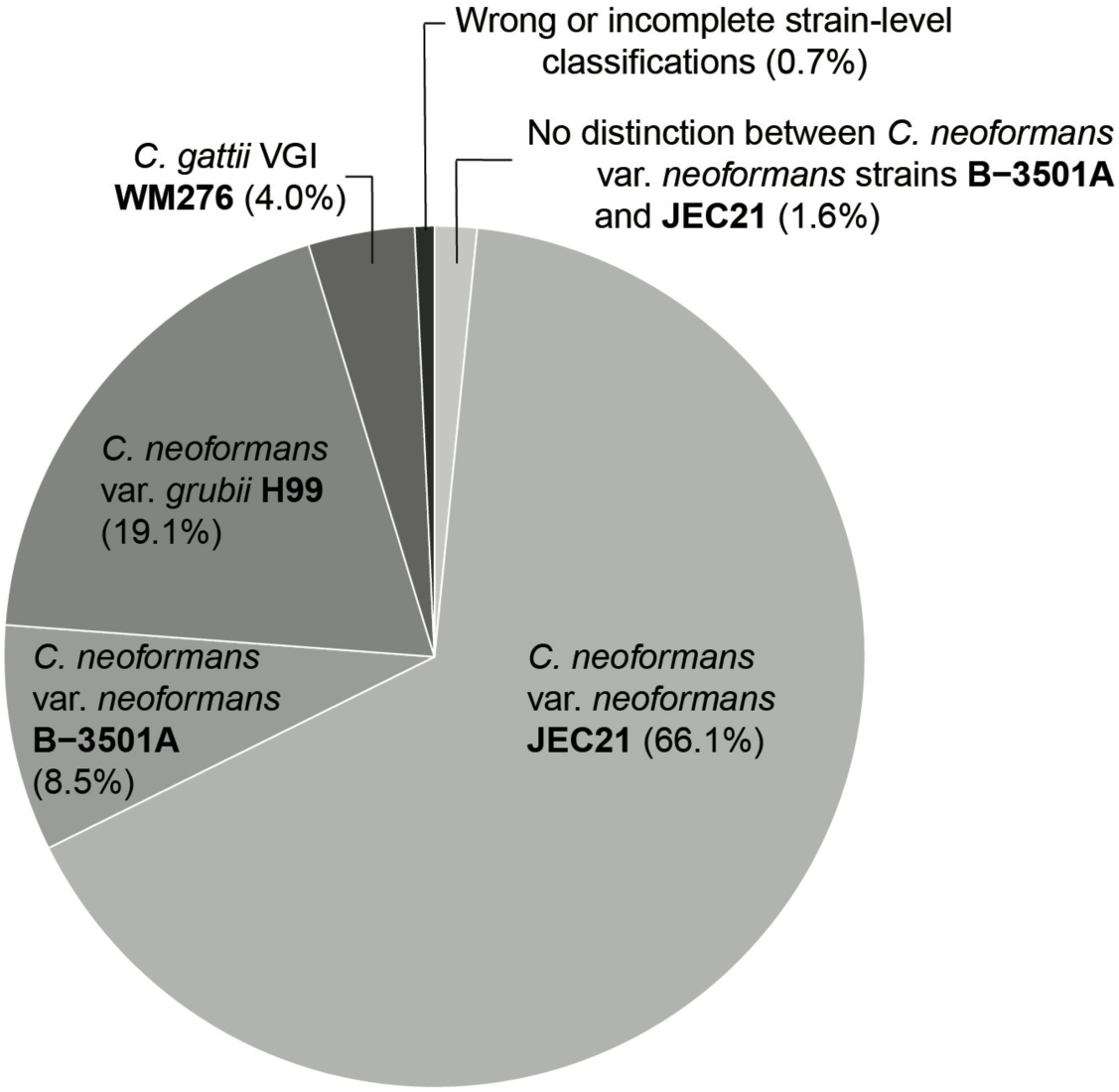
Strain-level classifications of taxa within the *Cryptococcus neoformans* and *C. gattii* species complexes.

## Discussion

Our metatranscriptomics approach yielded taxonomic identification of fungi from a defined mock community with high success, while false-positives were detected at far lower abundance. These results indicate that it is possible to obtain accurate species- and strain-level identifications for fungi from metatranscriptome data, as long as taxa identified at low abundance are removed from the analyses to avoid false-positives derived from contamination or misclassifications.

Taxonomic classification at species and strain levels using metabarcoding and metagenomic data has been considered inaccurate (Nilsson *et al.* 2019; Sczyrba *et al.* 2017; Yamamoto *et al.* 2014), raising the question of how our metatranscriptomics approach succeeded in identifying closely-related fungal strains. Metabarcoding relies on a single marker gene (Banchi *et al.* 2018; e.g. McGuire *et al.* 2013; Schmidt *et al.* 2013), which does not contain sufficient phylogenetic information to differentiate some closely related fungal lineages (Balasundaram *et al.* 2015; Nilsson *et al.* 2008). Metatranscriptomics, on the other hand, yields data on all expressed coding sequences. Classifications derived from metagenomes are likely to be equally accurate as the ones obtained from metatranscriptomes, except that dead organisms might also be sequenced. Additionally, we used a new alignment method that is both highly accurate and fast (Clausen *et al.* 2018), allowing us to map sequences against the complete NCBI nucleotide collection. Complete genomes of all fungal isolates used here are available in NCBI, and it is likely that the accuracy of identifications is reduced for poorly-documented microorganisms. However, it is possible to extract informative genes from metatranscriptome data and subsequently perform phylogenomic analyses to identify rare and novel taxa (e.g. Shi *et al.* 2017; Wille *et al.* 2018; Zhang *et al.* 2018). Besides being highly accurate, metatranscriptomics is less susceptible to amplification bias, no information about the community members is required *a priori*, and it only detects functionally active members of a microbial community. These advantages make metatranscriptomics a promising tool in biodiversity surveys, functional assessments of microbial communities, pathogen detection and biosecurity surveillance (e.g. Kuske *et al.* 2015; Shi *et al.* 2016; Wille *et al.* 2018).

Even though false-positives were present at low abundance, they pose a challenge in the interpretation of metatranscriptomic and metagenomic data. False-positives generally result from spurious classifications and laboratory contaminants, which may be common in laboratory reagents (Salter *et al.* 2014). However, metatranscriptomics is less sensitive to laboratory contamination than DNA-based metagenomics or metabarcoding, as only living microorganisms are sequenced. Nevertheless, contamination can occur at all stages of the library preparation and is routinely observed in RNA-Seq studies (Quince *et al.* 2017; Strong *et al.* 2014). Misclassifications occur because some genome regions are very similar (or identical) across closely-related species and cannot be differentiated. Errors in reference databases can also result in misclassifications. Sequences attributed to incorrectly-classified species are not uncommon in GenBank and result in downstream classification errors (Li *et al.* 2018). It is also not unusual to find bacterial regions misassembled into eukaryotic genomes (e.g. Koutsovoulos *et al.* 2016), which can result in sequences from common laboratory contaminants being classified as a eukaryote. Filtering out organisms found in low abundance is an option to reduce the incidence of false-positives in downstream analyses. In this study, filtering organisms for which the abundance of transcripts is lower than 0.1% would eliminate false-positives, at the cost of excluding one true-positive from the analyses (Table 2). The application of this abundance-filtering step might not be feasible when sequencing depth (per microbial species) is limited. Species present in low abundances will be represented by a small number of transcripts and so are more likely to be misclassified or undetected.

The abundance of transcripts and sequence reads can vary according to genome size, number of coding sequences and gene expression. Therefore, the abundance disparity across species observed here is unsurprising. Interestingly, we found no correlation between the abundance of transcripts and genome size or number of genes (Supplementary figure 1). Imprecise estimates of cell abundance and RNA extraction biases could also have influenced abundance estimates, and might be the reason why two species in the mock community (*D. hansenii* and *S. pombe*) were not detected in the analyses. Metabarcoding studies have suggested that performing DNA extraction in triplicate minimizes biases for bacteria, but it had no effect in fungal communities (Feinstein *et al.* 2009). To our knowledge, the effect of RNA extraction bias in metatranscriptomics has yet to be studied. As metagenomics surveys are not affected by gene expression, they might be more appropriate for studies where it is important to quantify species abundance.

Although fungal species and their genes can be confidently identified, it remains challenging to link some genes with particular species using metatranscriptomics. A large portion of fungal genomes are highly similar among species, making it difficult, if not impossible, to infer which species in the community are expressing which genes. Recently, a method was developed to perform species-level functional profiling of metagenome data (Franzosa *et al.* 2018). This method, however, relies on a reference database of complete genomes that currently contains few fungal representatives, limiting its application in fungal metagenomics. Contrary to metatranscriptomics, metagenomics yields coding and non-coding sequences, which can facilitate linking genes to species if sequencing depth is large enough to assemble large parts of fungal genomes (e.g. Olm *et al.* 2019).

In sum, we show that metatranscriptomics is a useful approach to identify fungal species and subspecies in mixed samples. The major advantages of metatranscriptomics over other HTS technologies include the selective sequencing of living organisms and the ability to detect a wide range of microorganisms in one step, which has multiple applications in biological research, surveillance and diagnosis. There is an increasing literature reporting that virulence and antimicrobial tolerance traits vary within species (Firacative *et al.* 2016; Rizzetto *et al.* 2013; Schauwvlieghe *et al.* 2017; Strope *et al.* 2015) and that multiple strains or species can co-infect a host (Desnos-Ollivier *et al.* 2010; Gupta *et al.* 2014; Seki *et al.* 2014; Soll *et al.* 1988; Tati *et al.* 2016). The high discriminatory power obtained for closely-related lineages of *Cryptococcus* provides a good example of where metatranscriptomics would be valuable in precision medicine, where therapy practices are defined according to strain-specific pathogenicity and drug susceptibility traits. However, it must be acknowledged that metatranscriptomics also has limitations that are common to high-throughput sequencing methods, as it is susceptible to DNA/RNA extraction biases, contamination and misclassifications. These limitations can be significantly minimized if appropriate controls are in place (*e.g.* abundance filtering before statistical analyses). Besides its application to identify well-known fungal species, metatranscriptomics can help to identify novel functional roles of fungi (e.g. Gonzalez *et al.* 2018; Liao *et al.* 2014) and novel species when used within a phylogenomic framework.

## Acknowledgments

We thank Fabio Santos and Krystyna Maszewska for helping with the fungal cultures and cell abundance estimates, and the High Performance Computing support at Sydney University. TCS and VRM are Sydney Medical Foundation Fellows whose work is supported by the Sydney Medical School Foundation. This work was funded in part by an NHMRC Centre of Research Excellence Grant (APP1102962) to TCS and an NHMRC grant (APP1121936) to WM and TC. ECH is funded by an ARC Australian Laureate Fellowship (FL170100022).

