## Supplementary figure for "Metatranscriptomics as a tool to identify fungal species and subspecies in mixed communities"

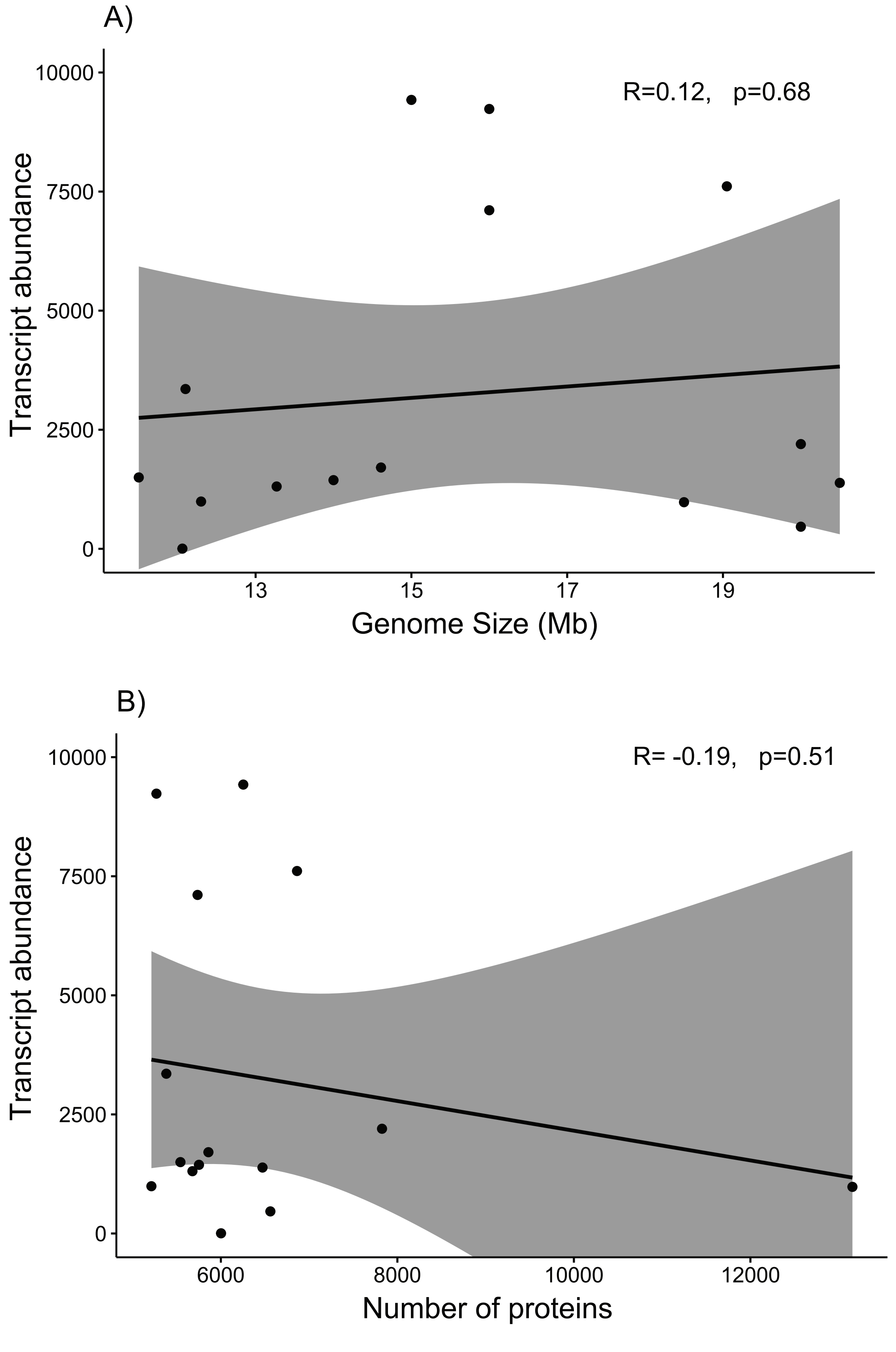


**Supplementary Figure S1.** No correleation between abundance of transcripts (depth) and genome size (A) or protein count (B) was observed. Each dot represents a fungal species included in the mock community. Pearson's correlation R and p values are given. Shaded areas indicate 95% confidence interval.
